# A new framework for analyzing mass transport in cortical brain tissue at <10 nm resolution

**DOI:** 10.1101/2025.04.02.646802

**Authors:** Donald L. Elbert, Dylan P. Esguerra

## Abstract

Transmission electron microscopy of brain tissue yields high resolution maps of the spatial organization of cells in the central nervous system. Automated segmentation identifies distinct biological cells and the resulting segmented images are easily converted into surface meshes. The surface meshes are structural models of cell surfaces, providing a framework for modeling of mass transport within the interstitial fluid of brain tissue that surrounds the cells. Our goal is to model the production and clearance of proteins implicated in the development of Alzheimer’s Disease. This work introduces a new custom computational framework to allow massive parallelization of the solution of the mathematical equations of mass transfer. The diffusion equation was solved directly on the surface meshes of multiple biological cells in parallel, with exchange of mass across co-localized faces at the end of each time step. Mass transfer across the interfaces of multiple analysis volumes was incorporated using a semi-implicit approach. To demonstrate the capabilities of the framework, unsteady mass transfer along cell surfaces was modeled in an array of eight 4 x 4 x 4 μm analysis volumes containing 2175 biological cells at 8 nm resolution, consisting of 313 million face elements. Areas of enhanced and hindered diffusive transport were identified, suggesting structural motifs that may contribute to the development of insoluble plaques. The tortuosity and fractional anisotropy were consistent with Diffusion Tensor Imaging (DTI) measurements in cortical tissue. This new framework allows modeling of diffusive mass transport while preserving anatomical details at <10 nm resolution.

**Author summary:** A framework was developed to model mass transport around thousands of biological cells in parallel. This may be useful to identify structural features in cortical tissue that are more prone to the development of amyloid plaques.

## Introduction

The automated serial processing of samples for high resolution transmission electron microscopy provides an unprecedented view of the architecture of the central nervous system. The Allen Institute in 2020 published an open dataset of aligned serial sections from 1 mm^3^ volume of mouse visual cortex.^1,2^ The resolution of the images was 4 nm x 4 nm in the x and y directions with section thicknesses of about 40 nm. Individual cells were identified within 8 nm x 8 nm voxels in the x and y directions and 40 nm in the z-direction. Automated segmentation algorithms were supplemented with manual corrections in some areas.^3–7^ Additionally, a 1 mm^3^ volume of human cortex has been released by Lichtman and colleagues.^8^

The rich dataset from mouse visual cortex has been analyzed with a focus on the relationship between structure and electrophysiology.^4,9^ Others have focused on the vasculature and the surface of the brain, which has yielded insights into the role of glymphatics in the clearance of solutes from the brain.^10,11^ Glymphatics is a process by which cerebrospinal fluid mixes with cortical interstitial fluid along the walls of penetrating blood vessels.^12^ This provides a mechanism by which harmful molecules such as amyloid beta and tau are cleared from the brain. Our own work suggests that about 20% of amyloid beta is cleared from the brain by glymphatics, 5% is cleared across the blood brain barrier, and about 70% is cleared by proteolysis within the brain tissue itself.^13^ However, there is still a great deal of uncertainty about the nature and magnitude of these processes. The questions are relevant to the development of therapeutics that affect proteolysis, glymphatics or blood-brain barrier, each of which has an unknown upper limit on the effectiveness of the intervention.^14–17^

Although the level of structural detail required to accurately model these processes is still unknown, models using the full resolution of the TEM images may provide important information that would be missed with coarse grained models. Specifically, measurement modalities such as Diffusion Tensor Imaging (DTI) observe structural barriers to diffusion at a resolution of about 1 mm^3^.^18^ This may miss certain structural features that are poorly cleared by convection or diffusion, providing a locally high concentration of toxic species that may promote the formation of insoluble aggregates.^19^

In the current work, we present a framework that models diffusion around each cell (and cell fragments) in eight 4 μm x 4 μm x 4 μm volumes of visual cortex in the Allen Institute’s mouse Microns dataset at the full 8 nm voxel resolution. This work builds on the prior analysis of convection and diffusion in a wide-variety of structure-based models that account for the geometry of brain tissue at varying scales.^20–26^ There are also many studies that utilize realistic 3D structural models of neurons from light microscopy, electron microscopy and the NeuroMorph repository to model electrophysiology or DTI.^9,27–31^

One of the major challenges of the analysis is the lack of information about the extracellular space in the TEM images. Collapse of highly hydrated extracellular matrix during processing of samples for TEM (i.e. dehydration) will reduce the apparent space between cells, although in some regions the presence of receptor-receptor interactions already limits the distance between cells. Co-registration of neuron somas between live calcium imaging and TEM revealed shrinkage in the x and z directions of less than 10% and expansion in the y direction of about 15%.^4^ While this leads to a negligible overall volume change, cryoelectron tomography (cryo-ET) images suggest a much more complex architecture of the extracellular space.^32–34^ TEM provides little information about the size and structure of the extracellular space, so relatively strong assumptions are required at the present time about the structure of the extracellular space throughout the cortex.

Even with perfect knowledge of the structure of the extracellular space, the high aspect ratios present computational challenges (e.g. space between cells may be as small as a few nanometers in synaptic clefts to 40-100 nm in other areas, while axons and dendrites can extend centimeters or more within the cortex). In this work, diffusion directly on surface meshes is modeled, reducing complexity but at the cost of accuracy for highly distorted meshes. Modeling diffusion along surface meshes has been used as a computationally efficient method to calculate geodesics between points on a complex surface mesh.^35–39^ Despite errors inherent in the present approach, it does provide a proof-of-concept and framework for nanoscale modeling of the cortex. The transition from the surface mesh approach to a 3D approach will be greatly aided by advancements in mesh smoothing algorithms. A novel smoothing algorithm used in this work has already greatly improved the quality of the mesh while maintaining interfaces between adjacent cells and is described in detail in a companion publication.

Traditional computational fluid dynamics packages (e.g. ANSYS, FreeFEM, OpenFOAM etc.) were not developed with the current problem in mind. Biologically focused packages such as Simvascular and Neuron are powerful suites of computational tools but not designed to parallelize the analysis of flow around thousands of unique objects.^40,41^ Therefore, we addressed this problem with a custom CFD framework. The segmented image stacks were smoothed and converted to surface meshes using a parallelized modification of the Lewiner Marching Cubes algorithm in C. From there, custom Julia code was used to record nodes, edges and faces for fast lookup, with most of the quantities needed for solving the diffusion problem pre-recorded. Data and algorithmic structures are inspired by OpenFOAM but specialized for the current problem.^42^

The Julia language was chosen because of the potential for C-like speed combined with ease of development, excellent support for simple memory management, linear algebra and parallelization on a high performance cluster (through MPI.jl).^43,44^ Furthermore, preliminary tests have indicated that there was little to no speed up by re-writing the code in C as long as type stability was ensured and allocations were minimized. Loading times (time to first ‘X’) are an issue with Julia and other non-compiled language, so large, general purpose scientific Julia packages (ODE/PDE packages, mesh and FEM frameworks, plotting tools, etc.) were avoided.

We will describe here the software framework and present result for an 8 μm x 8 μm x 8 μm portion of the visual cortex to illustrate our approach to encoding the geometry, properties and boundary conditions and parallelization of the analysis.

## Results

As a first step toward modeling pressure-driven glymphatic flows, we investigated diffusion around cortical cells through the extracellular space. Representative TEM images and surface meshes from the

MicronsExplorer website reveal how tightly packed cells are in the cortical tissue, which is due at least in part to dehydration of the tissue during sample preparation (Figure 1A). The region of interest is located in layer 3, which is a site of early development of Alzheimer’s plaques in humans as well as mouse models of the disease.^45,46^

**Figure 1:**
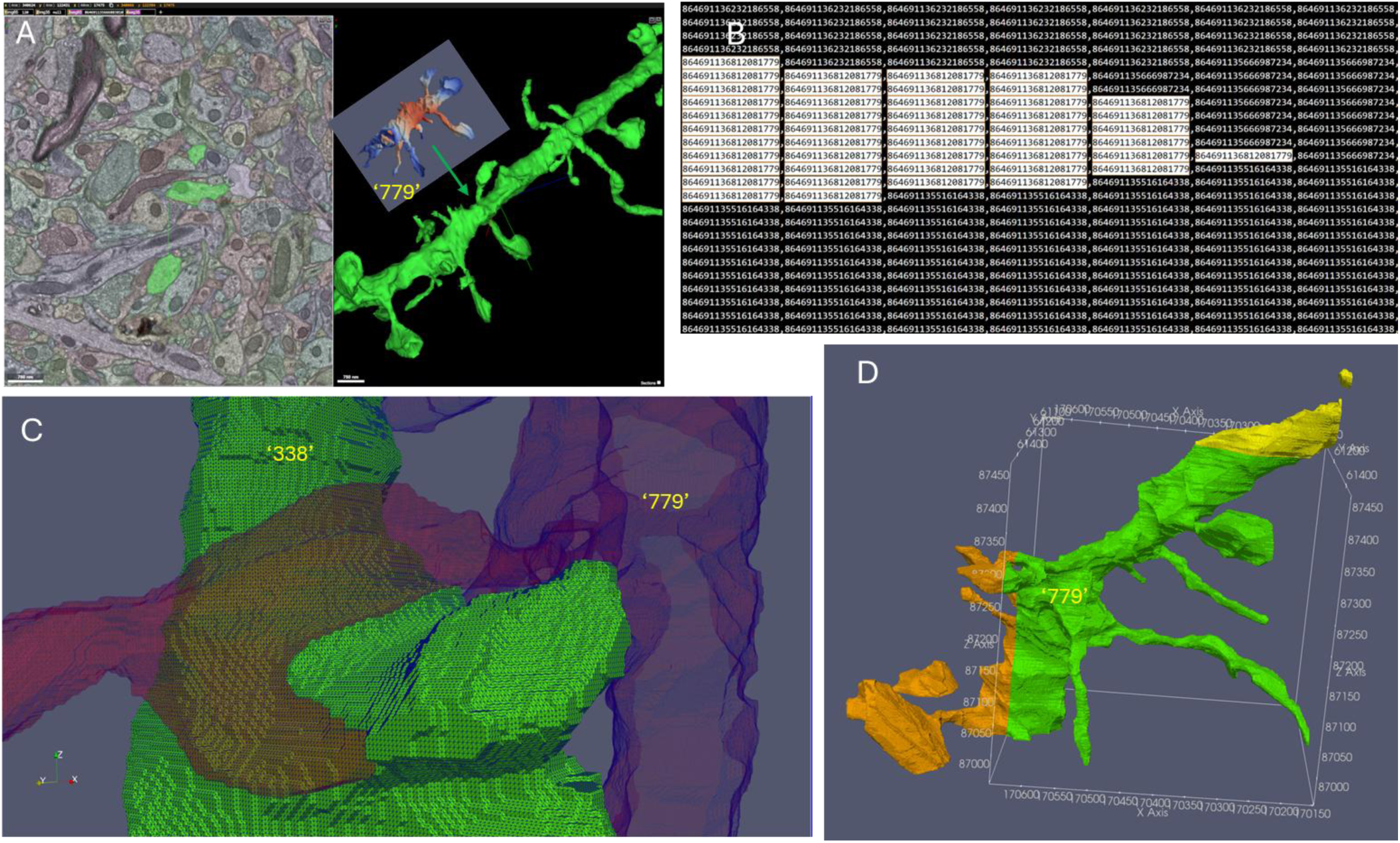
A. Allen Institute TEM data from mouse visual cortex (left), and a surface mesh of cell 864691136812081779 (‘779’; right), visualized on the MICrONS Explorer website (https://www.microns-explorer.org/cortical-mm3). Scale bars are 750 nm. B. The raw segmented data from a slice in the XY plane. Voxel size is the segmented dataset is 8 x 8 x 40 nm. Each voxel records an 18 digit identifier to a unique biological cell. C. The Lewiner Marching Cubes algorithm was applied to the segmented z-stacks downloaded from the MicronsExplorer database after mode filtering and image morphing to reduce z layer thickness to 8 nm. The algorithm results in defect-free meshes with perfect matching between faces on adjacent biological cells. ‘338’ = 864691135516164338 D. Cell ‘779’ within a 4 x 4 x 4 μm analysis volume and extending into adjacent analysis volumes.

The cell labeled in green is a layer 2/3 excitatory pyramidal neuron and the surface mesh is a branch of the apical dendrite. All cells in the dataset are labeled with an 18-digit number, and this cell is 864691136812081779 (‘779’). This region was chosen because it is near the center of a capillary bed, is morphologically consistent with a layer 3 neuron, and cell ‘779’ is one of five cells displayed at the launch of MicronsExplorer cortical-mm3 in Neuroglancer (https://www.microns-explorer.org/).

The surface meshes in MicronsExplorer and the false coloring of the TEM images shown in Figure 1A are the result of a segmentation algorithm. While one columnar area in the dataset has been extensively reviewed by humans, the current region of interest mostly reflects the automated segmentation algorithms.^7^ The methods shown here thus will be amenable to any segmented image stack even without human validation. The segmented images are two-dimensional arrays of 18-digit identifiers (Figure 1B). While images are at 4 nm resolution, segmentations are at 8 nm resolution. Figure 1B shows cell ‘779’ highlighted in the segmented image surrounded by two other cells. The segmentations that were used were version v343, although the methods were also tested with v943. Recently, version v1300 was released. Other than changes to some of the 18-digit identifiers, very few substantive changes were observed between v343 and v943 in this region.

While surface meshes are available from the MicronsExplorer website, the segmentations contain some obvious errors (isolated voxels) that are biophysically impossible (too small to accommodate bending of the plasma membrane). These were removed by mode filtering, with a few added rules to handle the presence of three or more cells in a single 3 x 3 window and rules to preserve corners of squares. The latter is important when 3 or more cells meet, as successive application of the mode filter may lead to erosion of the cell starting from square corners. Another rule that was applied was to replace cell assignments if a cell was represented in the inner 10 × 10 area of an 11 x 11 window but was not represented on the edge of the window. The application of these smoothing rules is described in a companion publication. After initial mode filtering, images are morphed (interpolated) in the z-direction using a new specialized method that preserves interfaces between cells and does not require identification and tracking of boundaries between the cells, described in the same companion publication. The difference in resolution in the z direction (nominally 40 nm) versus the x and y directions (8 nm) could be addressed by downsampling in the x and y directions although fine structures such as dendritic spines are greatly affected by such downsampling (data not shown). Instead, 8 nm resolution in the z-direction was achieved by morphing between frames over five layers, four of which are interpolated. This is followed by additional mode filtering. Representative surface meshes after mode filtering and morphing are shown in Figure 1C. Finally, surface meshes are contained within discrete 4 μm x 4 μm x 4 μm ‘analysis volumes’. Multiple volumes may be analyzed simultaneously if mass is allowed to transfer across the analysis volume faces at each time step. Extension of cell ‘779’ across multiple analysis volumes is shown in Figure 1D.

The modeled volume required to capture a capillary bed will be between 32 μm x 32 μm x 32 μm and 128 μm x 128 μm x 128 μm. The center of the image in Figure 2A is the center of the starting 4 μm x 4 μm x 4 μm volume. At a similar resolution, Figure 2B shows the somas of three cells that are in contact within the starting volume, cells ‘779’, ‘338’ and ‘435’ (‘338’ = 864691135516164338; ‘435’ = 864691136334080435), showing that their somas are well within the 128 μm x 128 μm x 128 μm volume. Cell ‘825’ (‘825’ = 864691136024224825) is a layer 6 neuron whose soma is well out of frame (approximately 450 μm distant).

**Figure 2:**
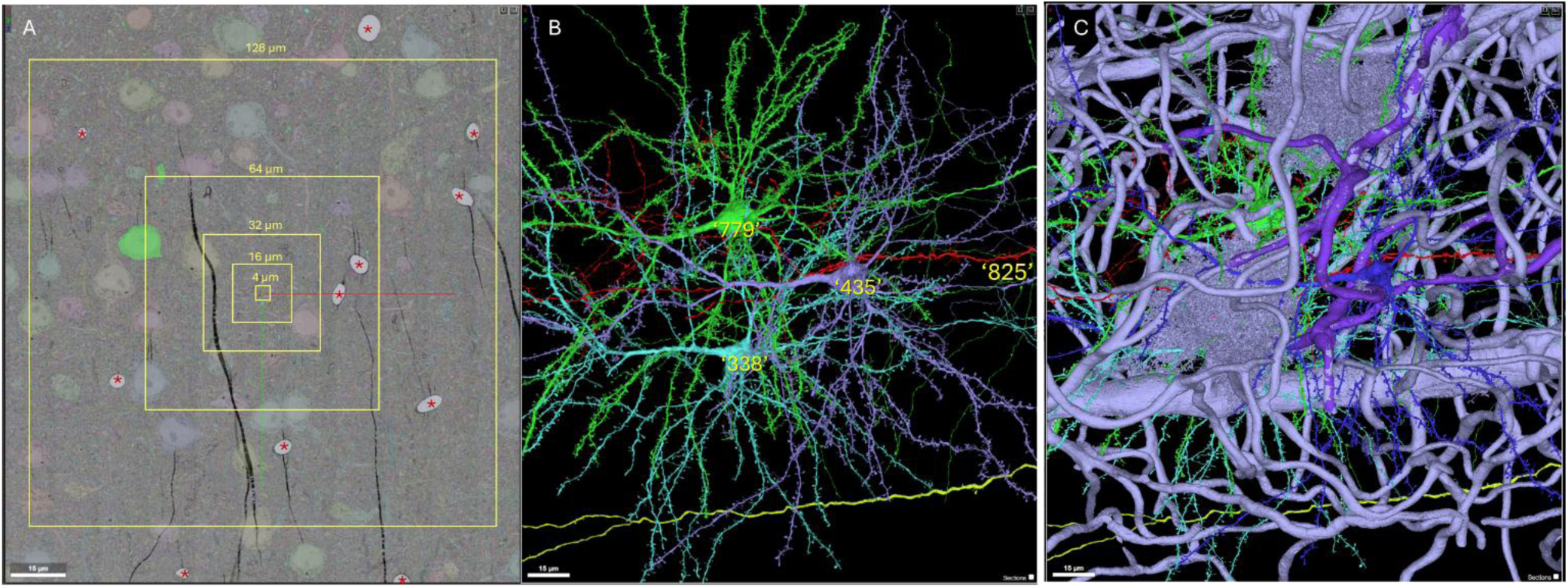
A. Visualizing the dimensions of a capillary bed in a section of mouse visual cortex (layer 2/3) on the MICrONS Explorer website (https://www.microns-explorer.org/cortical-mm3). Capillaries labeled with *. B. Surface meshes of selected neurons in this region. Cells: ‘779’ = 864691136812081779, ‘435’ = 864691136334080435, ‘825’ = 864691136024224825, ‘338’ = 864691135516164338. C. The capillary bed in this volume. Note that astrocytes are identified as being part of adjacent capillaries by the segmentation algorithm in the public dataset. Scale bars are 15 μm.

Although only a few capillaries are visible in Figure 2A, Figure 2C shows that the volume of interest contains numerous capillaries and some arterioles/venules.

The workflow for the analysis is shown in Figure 3. The mode filtered and image morphed segmented images are converted to Ply surface meshes using the Lewiner marching cubes algorithm using a triangular mesh over a grid with voxels spaced at 8 nm in all directions. The code was originally written in C++ by Lewiner and converted to C by others.^47^ The existing C code (https://github.com/neurolabusc/LewinerMarchingCubes) was parallelized by us to create surface meshes for multiple biological cells simultaneously using the Message Passing Interface (MPI). The marching cubes algorithm is applied to all cells within a 4 μm x 4 μm x 4 μm analysis volume. Although subsequent steps are amenable to using larger analysis volumes, the marching cubes algorithm as currently written is memory intensive and currently limits the size of analysis volumes.

**Figure 3:**
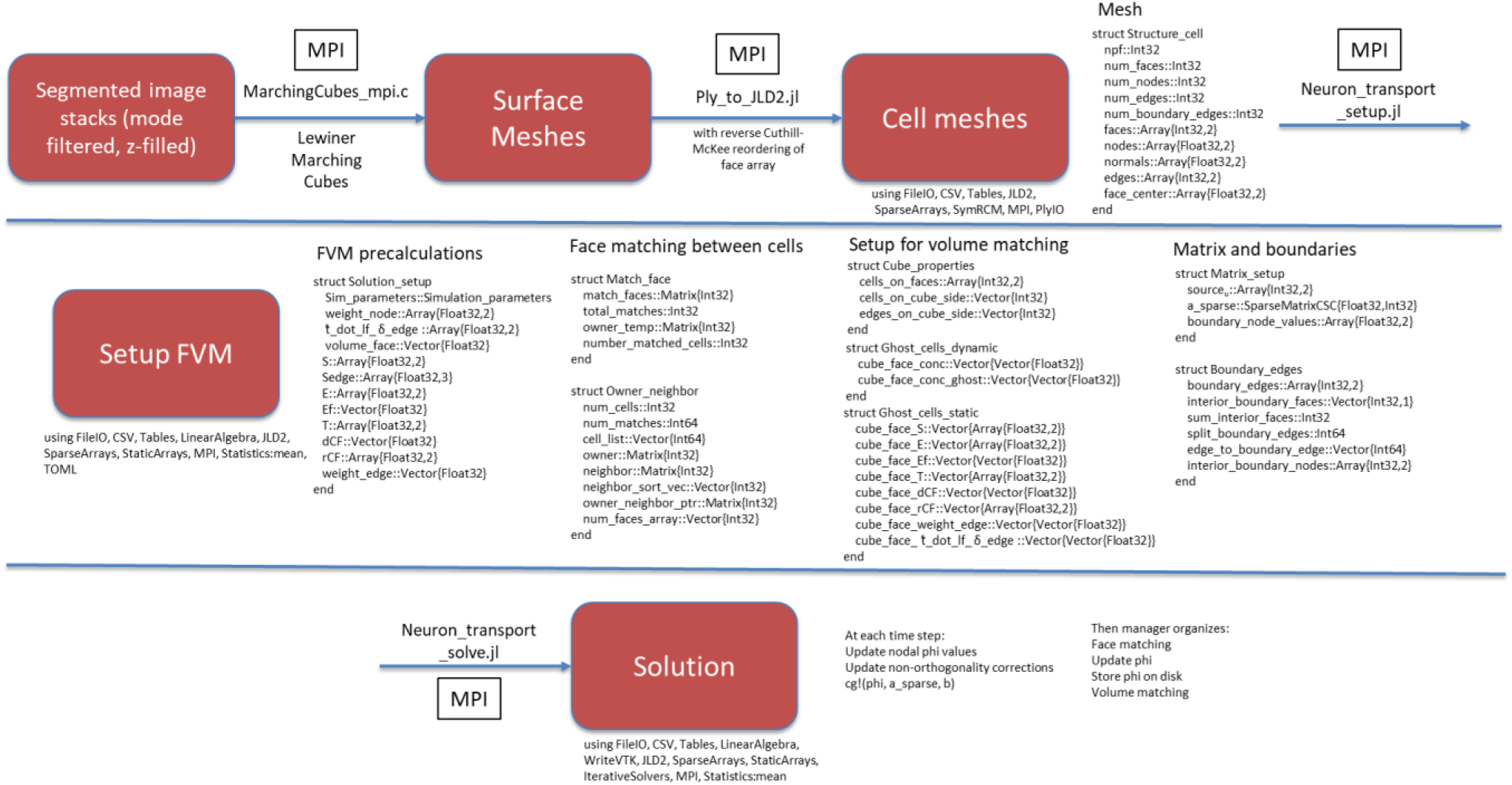
Workflow for modeling extracellular diffusion in brain cortex. Segmented image stacks were downloaded from the MicronsExplorer website. Following mode filtering and image morphing, the segmented voxels were converted to 3D surface meshes by the Lewiner Marching Cubes algorithm, with surface meshes stored as Ply files. The Ply surface meshes were analyzed and the resulting data were stored in the Julia JLD2 implementation of the HDF5 format. Pre-calculations were performed for: 1) finite volume analysis, 2) mass exchange between meshes that share faces, 3) mass exchange across edges on boundaries. Solution of the unsteady partial differential equations was performed in a parallel manner.

The resulting ply files are then analyzed to extract information about mesh edges and the data are stored in a single structure that is then compressed using a Julia-specific version of the HDF5 protocol called JLD2. All subsequent programs were written in Julia. Julia has an ease of use that is greater than C, with a sensible package management system and C-like multithreading/MPI capabilities. Julia may be viewed as a rapid prototyping language^48^ and has been used by Meta in this way (https://www.youtube.com/watch?v=3ypsZUNRjI4&t=493s).

After extraction of information about the meshes, a setup program discretizes the equations, handles boundary conditions and boundary edges, identifies mesh faces that are shared by separate biological cells (‘face matching algorithm’) and prepares for communication across the faces within an array of analysis volumes. The finite volume discretized equations are solved at fixed time steps by the implicit (backwards) Euler method. At each time step, the equations are solved for individual cells in the volume in a parallel manner using a manager/worker job scheduler. At the end of each time step, concentrations are averaged across matching faces and ‘ghost’ concentrations are stored at faces of cells that pass through the face of an analysis volume. The ghost concentrations are used during the next time step to estimate flux across the faces of the analysis volumes in a semi-implicit manner. Because of the need to exchange information across analysis volumes at each time step, the current rate limiting step of the analysis is the substantial overhead with loading and unloading the properties of each analysis volumes at each time step. Additional details may be found in the companion publication focused on the computational methods.

The computational framework was applied to all the biological cells in a single 4 μm x 4 μm x 4 μm analysis volume (Figure 4). The volume contained parts of 431 cells (Figure 4A). The cells are densely packed, five cells were selected for visualization, with four of which are neurons highlighted in Figure 2 and the fifth being an oligodendrocyte (Figure 4B). Because of our interest in modeling clearance of toxic species, for the model the initial concentration was 1.0 (arbitrary units) on all faces of the surface meshes of all 431 cells. For boundary conditions, the nodes on the faces of the analysis volume were fixed at concentration = 0.0. The concentration profile for all 431 cells is shown in Figure 4C at time = 760 μs. Figure 4D shows the five selected cells from panel B to illustrate the concentration profile in the interior of the analysis volume. The inset shows that the termini of the spines of cell ‘779’ are losing mass through interaction with neighboring cells. This illustrates the communication between cells via the face matching algorithm that mimics the presence of a small channel between adjacent cells.

**Figure 4:**
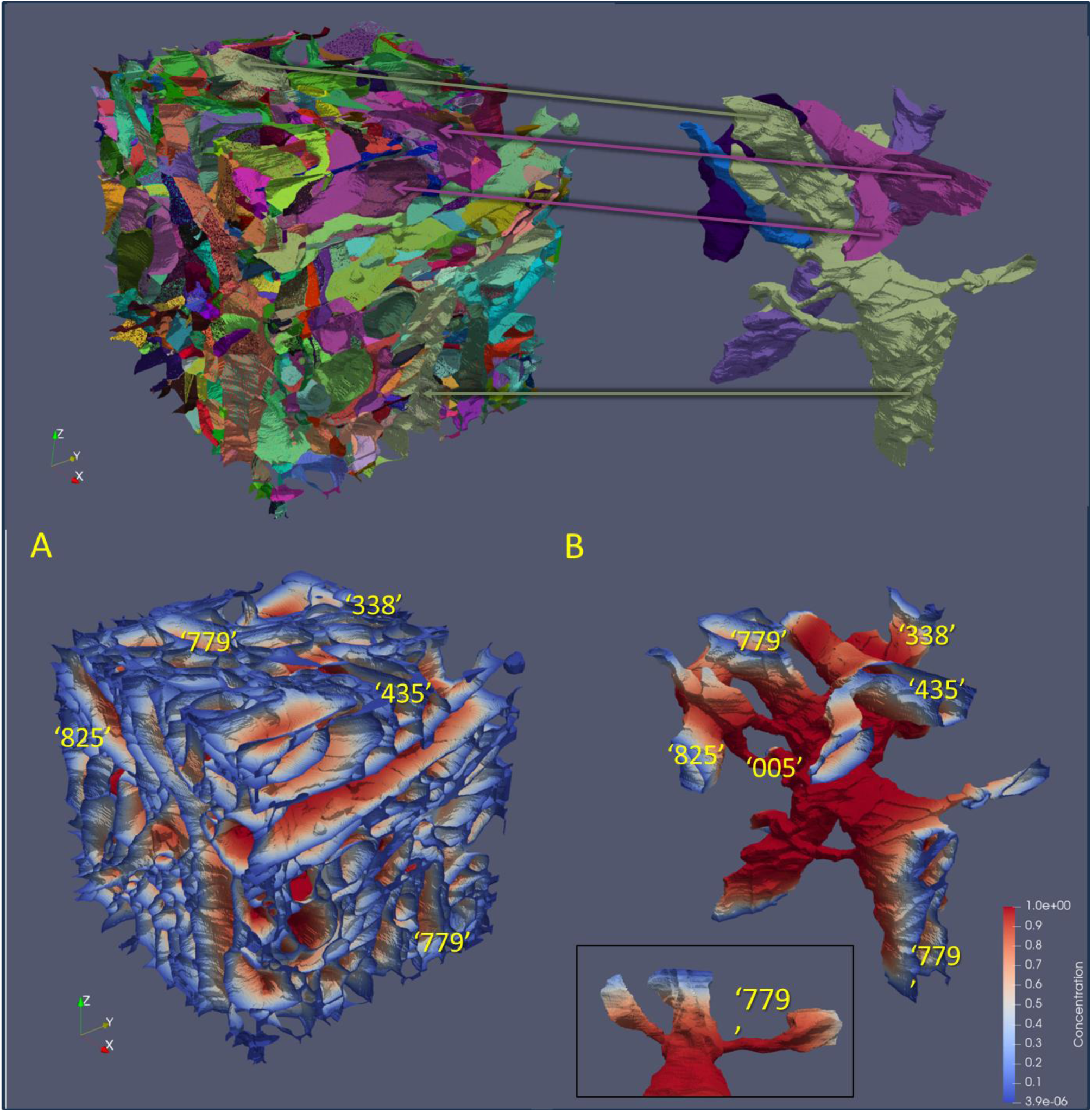
A. The 431 biological cells passing through a 4 μm x 4 μm x 4 μm analysis volume with center point at x = 170398, y = 61241, z = 17451 in ‘8 nm’ coordinates. B. For visualization, five cells are selected and shown, with arrows pointing to their location within the full set of cells. C. Solution for all 431 cells in the volume at time = 760 us. Initial condition: concentration = 1.0, concentration at boundaries = 0.0. The diffusion coefficient of the solute is 6.4 × 10-7 cm2/s, typical for a protein. Uniform time steps of 10 μs were utilized. D. Five cells from panel C, cells: 864691136812081779, 864691136662315005, 864691136334080435, 864691136024224825, 864691135516164338. The inset shows cell ‘779’ alone. The termini of the spines on ‘779’ are not connected to the surface of the analysis volume but show mass loss consistent with their distance to the boundary, illustrating the transfer of mass via the face matching algorithm.

Arrays of analysis volumes allow modeling of larger volumes of brain tissue. This is shown conceptually in Figure 5. A 2 x 2 x 2 array of 4 μm x 4 μm x 4 μm analysis volumes contain portions of about 400-500 biological cells that are in intimate contact with many other biological cells in the volume. The discretized equations are solved for each analysis volume sequentially, with ‘ghost’ concentrations at interior boundaries exchanged at the end of each time step. The ghost concentrations are used in the next time step to calculate fluxes across the boundaries in a semi-implicit manner. Independently, mass is also exchanged between neighboring cells within an analysis volume at the end of each time step. This mass exchange is in the form of averaging the concentrations at shared faces.

**Figure 5:**
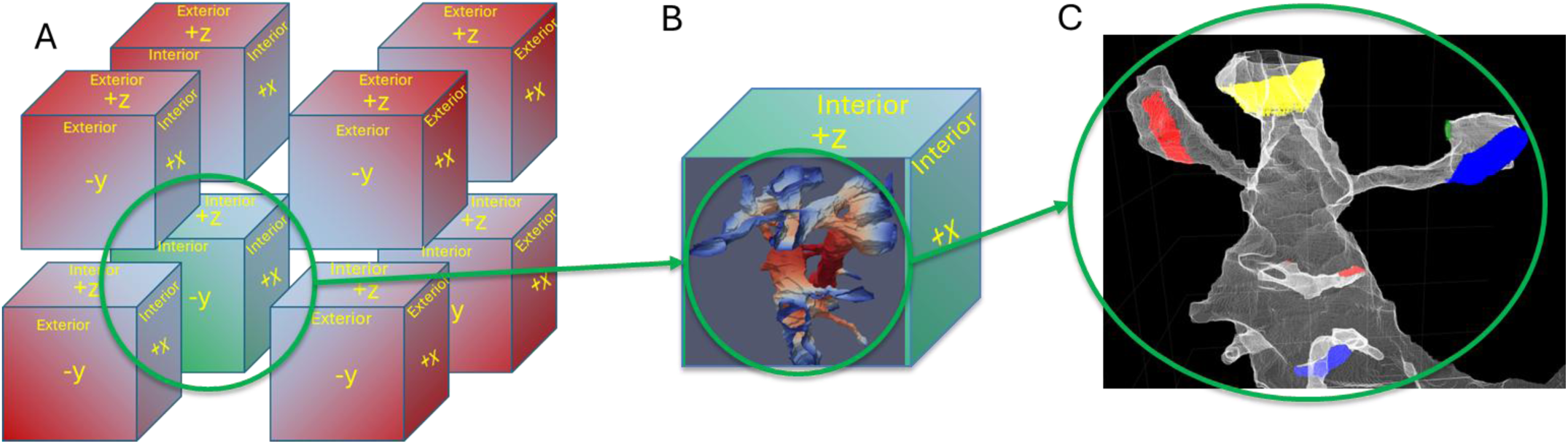
Arrays of analysis volumes. A. An array of eight 4 μm x 4 μm x 4 μm analysis volumes, with the volume from Figure 4 highlighted in green. B. Within each volume reside about 400-500 cell objects, which generally are much larger than the analysis volume and cross into neighboring volumes. The specified boundary conditions are applied at the faces on the exterior of the array, while mass transfer across interior faces requires exchange of data between the analysis volumes at each time step. C. Additionally, each cell in a volume is in contact with many other cells. Because the discretized equations are solved for each biological cell in the analysis volume independently and in parallel, mass exchange between neighboring cells must be performed at the end of each time step. The contact regions between the cell in white and four other cells are shown as red, yellow, blue and green patches.

Figure 6A shows results for an array of eight 4 μm x 4 μm x 4 μm analysis volumes, with 18 representative neurons shown. Solution of the equations for all eight analysis volumes was about 4.7 h per 1 ms of simulated time on a single node with 40 cores and 742 GB of memory. The number of nodes could be increased about 10-12 in the current configuration (400-480 cores) with close to a linear speedup as all cells in an analysis volume are processed simultaneously. After that point, the code would need to implement parallel handling of analysis volumes. However, with that addition, the array of analysis volumes could be arbitrarily large. The mean number of unique biological cells per volume was about 434 ± 31. As the volume doubled in size, the number of unique cells decreased by about 40%. For example, 2175 unique cells were present in the analyzed 8 μm x 8 μm x 8 μm volume while the total number of cell fragments in all analysis volumes was 3476. The total number of face elements in all meshes was 313,089,506. The inset to Figure 6A shows the eigenvectors of the effective diffusion tensor calculated from the data in Figure 7. The visible orientation of many neurons in the y-direction in Figure 6A, which is perpendicular to the surface of the brain is consistent with layer 2/3 architecture. The eigenvectors are oriented primarily along the cortical columns. Figure 6B shows the same 18 neurons, but at the first time step (thus mostly red with blue at the boundary,) with all of the astrocytes in the analysis volume (green). The astrocytes do not appear to have a preferred orientation.

**Figure 6:**
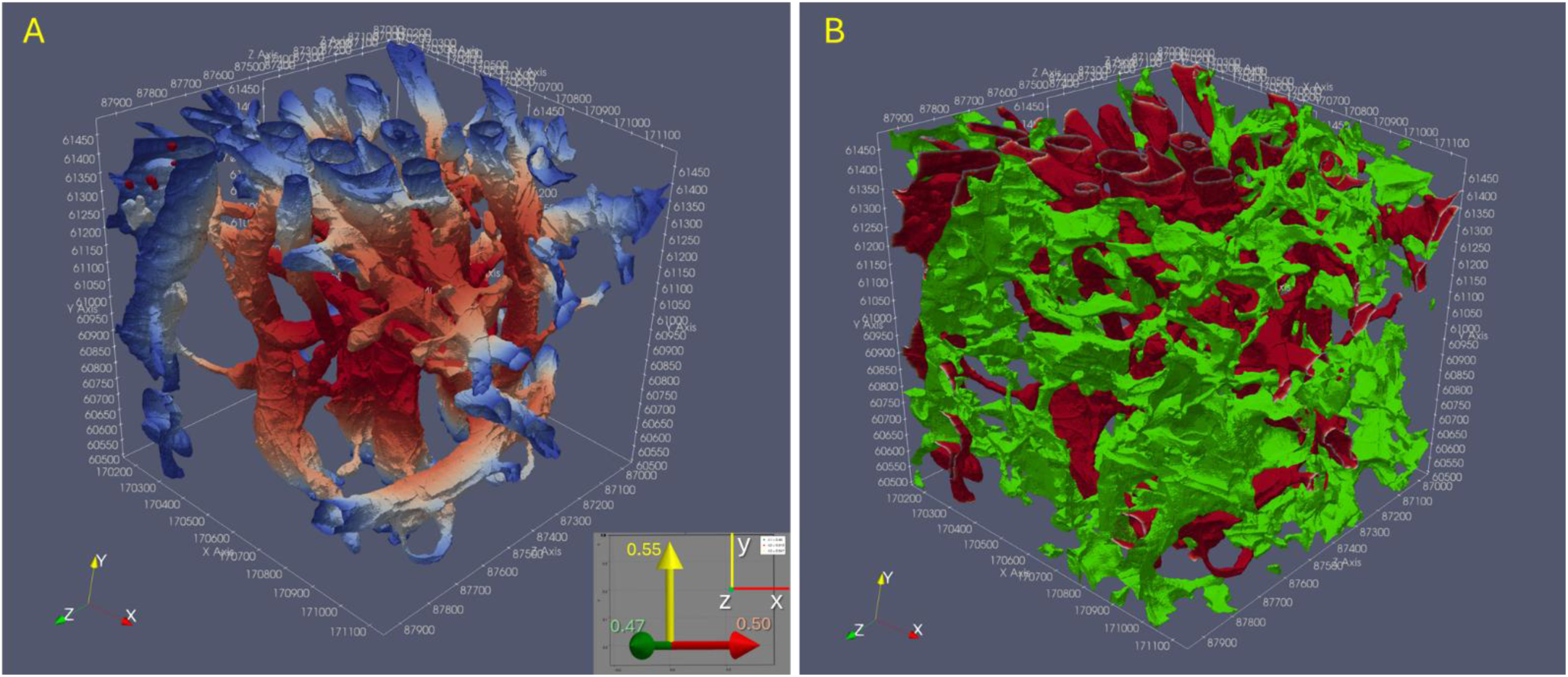
A. Eighteen neurons from an 8 μm x 8 μm x 8 μm volume that is an array of eight 4 μm x 4 μm x 4 μm volumes at time = 12.51 ms. Center point is x = 170648, y = 60991, z = 17491 in ‘8 nm’ coordinates. Inset are eigenvectors of the diffusion tensor (from Figure 7) viewed along the spatial z-axis. B. The neurons from part A at time = 1 μs (red except at boundary) shown with the astrocytes in the array of analysis volumes (green).

**Figure 7:**
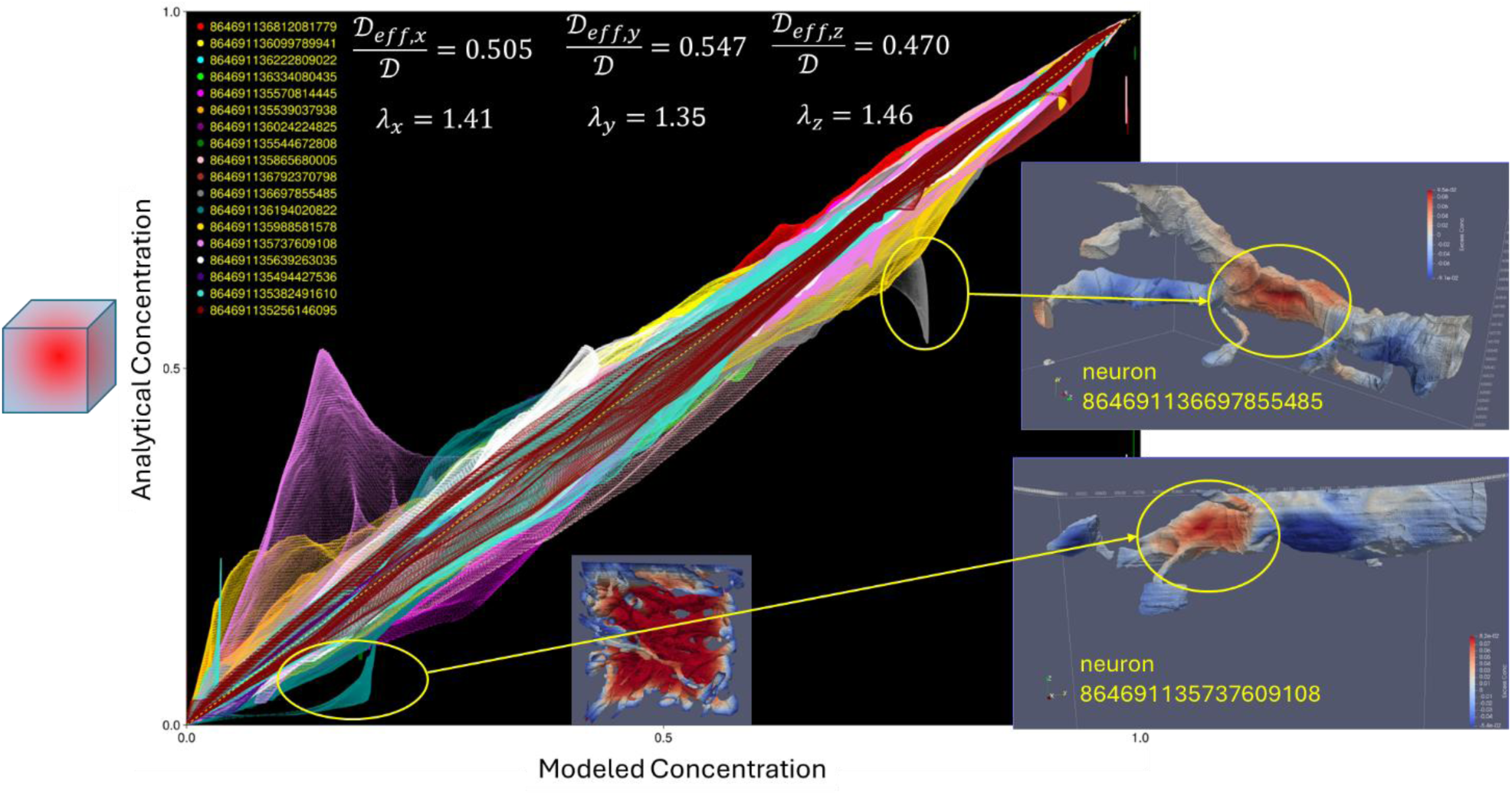
The concentrations of each face in the eighteen neurons from figure 6A at 25 ms (‘Modeled Concentration’) were compared to the predicted concentrations (‘Analytical Concentration’) for a solid cube of the same dimensions. The ‘effective’ diffusion coefficient tensor used to calculate the ‘Analytical Concentration’ was varied to minimize the difference between ‘Analytical’ and ‘Modeled’ concentrations. The ratio of the optimized ‘effective’ diffusion coefficient is shown for the x, y and z directions. The tortuosity was then calculated as the square root of the inverse of the diffusion coefficient ratio. Circled regions show slower diffusion (higher than expected concentration) that were observed on two neurons (insets). These ‘hot spots’ were associated with deep grooves opposite external faces, requiring diffusion out of the grooves prior to diffusion towards the boundaries.

For the 18 neurons in the eight 4 μm x 4 μm x 4 μm volumes, the concentration on each face of the surface mesh was compared to the expected concentration for diffusion from a solid cube with the same size, initial concentration and boundary conditions (Figure 7). The diffusion coefficient tensor in the solid cube was optimized to minimize the difference between the solid cube and simulated brain region. It was found that the y-component of the diffusion coefficient tensor was 8% higher than the x-component and 16% higher than the z-component. These are small differences compared to white matter, but reasonable for grey matter. The effective diffusion coefficient was about 0.5, with tortuosity of about 1.41. This is slightly lower than typical values measured by DTI, which may be due to the effects of extracellular matrix porosity that were not considered here and the local density of astrocytes in this volume (Figure 6D).^49^ The off-diagonal elements of the diffusion tensor were close to zero (<1×10^−9^) in the x- and z-directions but 0.02 in the y-direction, yielding a fractional anisotropy of 0.076, consistent with the architecture of grey matter being close to isotropic.

Any deviations from the line of identity (dashed yellow line) in Figure 7 indicate structures on the surface of the biological cells that have anomalously fast (above the line) or slow (below the line) diffusion. Investigation of some of these structures (yellow circles) revealed hot spots (red) in the excess concentration, indicating slower diffusion. These two examples were associated with deep grooves in the neuron plasma membrane that were due to the presence of cells such as microglia or oligodendrocytes. The deep grooves are shielded from the boundary by the cell itself, such that mass must diffuse in the opposite direction before diffusing towards the boundary. The presence of these structures may be sites of enhanced aggregation of amyloid beta, which is a testable hypothesis if TEM reconstruction was applied to mice that develop amyloid plaques. Knowledge of the existence of such sites would be helpful in predicting the distribution and timing of amyloid plaque accumulation with age, or re-emergence of plaques following removal with monoclonal antibodies. Addressing such questions is a primary motivation for development of the current modeling framework.

The structure associated with the most prominent peak above the yellow dashed line in Figure 7 is further investigated in Figure 8. This figure shows the same analysis as in the previous figure but for a single astrocyte (8646911356666842) instead of the 18 neurons. The same ‘cold’ or fast-diffusing area is seen at the same location as in Figure 7. The circled peak is due to the interaction of this astrocyte with neuron 864691135737609108 (see inset). The site of enhanced clearance may be due to the capping of part of the neuron by the astrocyte, providing unbranching pathways to the boundary. The astrocyte itself is essentially isotropic with the diagonal components of the effective diffusion coefficient at about 0.5 with a fractional anisotropy of 0.012. Even within this single nearly isotropic astrocyte, there are some structures (hot spots) that are poorly connected to the boundary.

**Figure 8:**
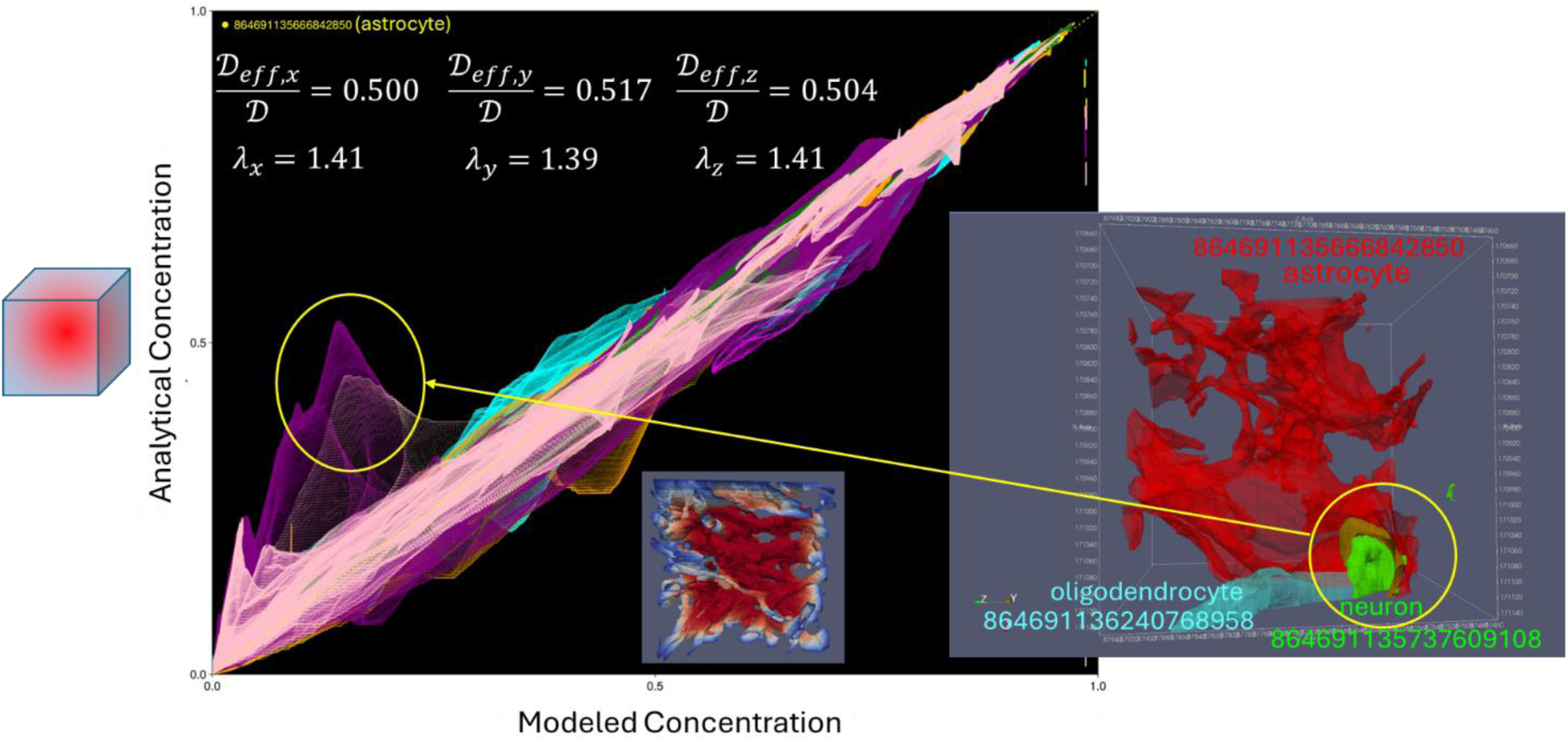
The concentrations (‘Modeled Concentration’) at 25 ms on each face of a single astrocyte (86491135666842850) were compared to the predicted concentrations (Analytical Concentration) for a solid cube of the same dimensions. The colors here represent the 8 different analysis volumes. The ‘effective’ diffusion coefficient tensor for the ‘Analytical Concentration’ was varied to minimize the difference between ‘Analytical’ and ‘Modeled’ concentration. The ratio of the optimized ‘effective’ diffusion coefficient is shown for the x, y and z directions. The tortuosity was then calculated as the square root of the inverse of the diffusion coefficient ratio. Circled region shows faster than expected diffusion (lower than expected concentration) that were associated with neuron 864691135737609108. This ‘cold spot’ was associated with the astrocyte completely covering the neuron and providing a route without branching to the boundary. An oligodendrocyte also helped provide a direct path to the boundary (864691136240768958).

## Discussion

Although the software used in the study is open-source, the focus of the current publication is on the overall framework for parallel analysis of diffusion in cortex at 8 nm resolution rather than a software product. The methods are under active development on several fronts, perhaps most importantly on producing smooth and biophysically accurate representations of the cells and their extracellular matrix. The meshes are most likely already smooth enough to successfully ‘inflate’ the surface meshes into 3D prism-based ‘inflation layers’ that would minimize the errors in the surface mesh-based solution of the diffusion equations. These inflation layers would likely need to be less 4 nm thick on each side to avoid elements with negative volumes, which is much thinner than the extracellular thickness. It is premature to produce inflation layers that are as thick as typical extracellular spaces. The average thickness of the extracellular space is difficult to measure but recent cryo-ET studies suggest close apposition of plasma membranes in synaptic clefts and other sites of direct receptor-receptor interaction, with lacunae that are 50-100 nm across.^32^ Cryo-ET reveals that the spines seem to have a narrower stalk than found by TEM and more spherical termini. The extracellular space between spherical termini results in wedge-shaped interfaces between cells. From a computational standpoint, wedge structures are unfavorable, leading to ‘pinched’ meshes. Spline-based meshes and phase field modeling may prove to be more efficient than traditional triangular/tetrahedral meshes.^50–57^ There are many open questions and opportunities for improvement of the framework.

Pressure-velocity coupling requires accurate calculation of gradients at faces, typically by the Green-Gauss method. For the current problem, gradients parallel to the edges were estimated via a nodal scheme that was used for a non-orthogonality correction via the overrelaxed approach.^42,58^ This is sufficient for diffusion but not pressure-velocity coupling. Thus, smoothing and inflation are areas of active development. In the current work, mass transfer by diffusion in eight analysis volumes was modeled (total volume = 8 μm x 8 μm x 8 μm). The optimal size of the analysis volume is currently unknown. With the current base volume of 4 μm x 4 μm x 4 μm, 32,768 analysis volumes would be required to reach our goal of 128 μm x 128 μm x 128 μm. Given 10 nodes with 40 cores each, the code should complete a time step in about 26 hours. On the one hand, it is impressive that the current analysis with 313 million elements is achievable at all in a reasonable time, while on the other hand the inflated mesh will increase the number of elements and pressure-velocity coupling will at least double the computation time. Improvements in speed and efficiency of the code, addition of pressure-velocity coupling and better modeling of the thickness of the extracellular space should be addressed before scaling the framework. Additionally, the exterior boundary of the analysis volume should be periodic while pressures and fluid flows should be specified at capillary surfaces. Additionally, while Julia is an excellent choice for rapid prototyping, there may be speed and memory improvements by implementing the methods in C.

Parallelization based on biological cells may be unnecessary. Fusing the cell meshes into a single mesh per analysis volume adds some complexity as the number of faces per edge will be >2 at interfaces between three or more cells. This may affect the stability of the solution methods and may be more prone to errors. In the current simulation, there are typically 0-2 cells per analysis volume that are small and near a boundary whose solutions diverge. These are handled in the current program by checking for NaN after solving the linear system and setting their concentration equal to zero if found. It is not yet clear if such domains would affect stability of the fused mesh. Additionally, keeping the cells separate may have advantages if the Hodkin-Huxley equations are solved in addition to pressure-velocity coupling and the mass transport equations. Another challenge is the dynamic nature of the cortex structure. Synapses may be pruned, axons and dendrites may shrink with sleep, blood vessels conduct pressure waves in their walls that lead to a pulsatile flow of interstitial fluid, and expression and distribution of aquaporin-4 in astrocytes may affect extracellular volume.

The face matching approach to mass transfer between adjacent surface meshes used here may present challenges if inflation layers are applied. It is possible to grow the meshes inward for each cell such that the equations are solved not at the surface mesh but within the inflation layer. If there is a one-to-one correspondence between the surface mesh and the inflated prismatic mesh, this would allow for the use of the same matching framework to be used. If the curvature of the meshes is too high and the thickness of extracellular matrix too great, then prismatic elements will need to be merged, breaking the connection to the surface mesh faces. Thus, advancements in the smoothing and inflation algorithms and the description of the extracellular space are crucial to future model development.

## Conclusion

The primary contribution of this study was to demonstrate the feasibility of handling thousands of objects across multiple analysis volumes. It may be difficult to adapt existing tools to this problem, which is the reason for the development of this independent framework. The primary insight is that some structural characteristics of cortical cells suggest potential sites of plaque deposition, which would not be seen in coarse-grained measurements or models.

## Methods

Simulations were performed on a 40 core Intel XEON 6230 Gold processor with 742 GB of memory. The Allen Institute MicronsExplorer dataset includes TEM images, segmentations and surface meshes. While surfaces meshes are available for download, we opted for greater control over the mesh generation process and started with stacks of segmented TEM images. The segmented TEM images contain 8 nm voxels, with each unique biological cell tagged with an 18-digit identifier. The distance between images in the z-direction is nominally 40 nm. To avoid skewness in the surface meshes, three approaches may be taken: 1) voxel size may be increased to 40 nm by downsampling, 2) the z-stack layers may be repeated five times at an 8 nm spacing, or 3) interpolations may be applied to add new frames to the z-stacks at an 8 nm spacing. We chose the last approach, and the interpolation scheme is described in a companion publication.

The location modeled within the TEM dataset has base imaging coordinates (x=340,796 y=122,482 z=17,391). This location was chosen because: 1) It is within cortical layer 3, a site of early plaque deposition in Alzheimer’s Disease, 2) near the center of a capillary bed, 3) Centered near a branch of the apical dendrite of one of the five cells shown at startup of MicronsExplorer. The listed x and y coordinates are at 4 nm resolution, while segmentation data is at 8 nm resolution. To convert from 4 nm coordinates to 8 nm coordinates, x and y coordinates are divided by two (x=170398, y=61241, z=17391). Starting from these coordinates, the appropriately sized slices were downloaded from: “precomputed://https://storage.googleapis.com/iarpa_microns/minnie/minnie65/seg_m343/”. The z coordinates are the same between the imaging and segmented datasets. However, because each z location represents a TEM slice that is 40 nm in thickness, a new coordinate system was used in which the z frame number is multiplied by 5 to avoid fractional frame numbers after insertion of four interpolated frames.

The Lewiner Marching Cubes algorithm was applied to smoothed and interpolated segmented image stacks. The original C++ version developed by Lewiner had been previously converted to C by others (https://github.com/neurolabusc/LewinerMarchingCubes). The C code was then modified here to add MPI capabilities, allowing processing of multiple biological cells concurrently.

The Lewiner Marching Cubes algorithm outputs PLY surface meshes for each biological cell within a given analysis volume. The PLY files (containing only nodes, face normals and face numbers) are further processed to extract information about nodes, edges and faces that is needed in the finite volume method. Nodes are stored in a 3 x *n_nodes* Float 32 array, which consist of node locations in analysis coordinates (*n_nodes* = number of nodes). Edges are stored in a 11 x *n_edge*s Int32 array, recording both nodes of an edge, boundary information if applicable, edge order in each face and adjacent faces (*n_edges* = number of edges). Faces are stored as a 12 x *n_faces* Int32 array, recording nodes, edges, adjacent faces and orientation of vectors joining face centers (*n_faces* = number of faces). Note that Julia arrays are column major and allow faster access to columns. The edge array is further processed to extract a ‘boundary_edges’ array for edges that lie on the face of an analysis volume.

Arrays of analysis volumes may be analyzed. In this paper, we begin with analysis volumes that are 4 μm x 4 μm x 4 μm in size. Eight of these volumes are then assembled in an array (8 μm x 8 μm x 8 μm). Analysis volumes are analyzed sequentially, with the flux across boundary edges incorporated semi-implicitly at each time point. Although there is no limit to the number of analysis volumes that may be analyzed, this number must be decided at the start of the simulation. The analysis volumes are not limited to 4 μm x 4 μm x 4 μm, but the memory requirements for the current marching cubes method become limiting as the volume grows. In the simulation settings, a ‘height’ is provided, which is the length of one side of the analysis cube, in um. The number of z-slices per μm is 25, and the number of voxels in the x and y directions is 62.5 per micron. To avoid fractional voxels, the ‘height’ in microns must be divisible by two.

The diffusion problem is solved for each biological cell in an analysis volume separately in parallel using MPI. The linear system to be solved is populated ahead of the simulation by the setup program, which also identifies matched faces of cells that pass through interior boundaries of analysis volumes. Boundary conditions are applied at the exterior boundaries of analysis volume arrays. At the end of each time step, the average concentrations of matched faces are recorded. Finally, calculation of the fluxes between adjacent analysis volumes is facilitated by pre-calculating the implicit part of the flux and setting up parameters for calculation of the explicit part. Space is pre-allocated on a manager core to allow storage of matched face concentrations and ghost concentrations, which is then shared with workers via one-sided communication.

After setup, the convection/diffusion equations are solved over time using uniform time steps. The diffusion coefficient for the solute was 6.4 × 10^−7^ cm^2^/s with time steps of 10.0 μs. To facilitate distributed analysis, solutions are stored on disk at each time step.

Simulations were performed in Julia 1.11.1 with the packages listed in Figure 3. Results were saved using the VTK format and visualized with Paraview 5.11.1. The exact solution for diffusion from a solid cube was (with N = 10, t = 25 ms):

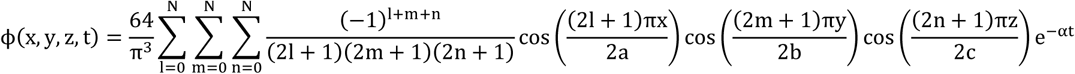

where

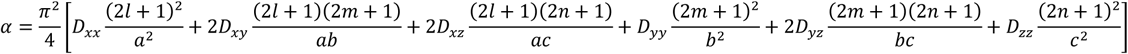

The excess concentration was the difference between the simulated concentration and the exact solution for the solid cube at the same location in space. The sum of squares of excess concentration was minimized by varying the six unique components of the diffusion tensor by the Nelder-Mead algorithm in Julia 1.11.1 with Optim v1.11.0. Plots were generated with Julia 1.11.2 and GLMakie v0.10.18.

## Acknowledgements

The work was supported by startup funds at the University of Washington

## Author contributions

DLE designed the study, designed the framework, wrote code and wrote the paper

DPE aided with design or the study, provided feedback on the framework, wrote code and edited the paper.

## Data availability

Programs used to generate the results are available at: https://github.com/elbert5770/neuron_transport_3D_v3_archive. Video results are available:

Figure 4: https://youtu.be/0ALJm-rfFp0

Figure 6: https://youtu.be/agXYVPlUfMc

The source data was downloaded from the Microns Explorer website: https://www.microns-explorer.org/cortical-mm3

